# Polyene-based colouration preserved in 12 million-year-old gastropod shells

**DOI:** 10.1101/2023.05.17.541181

**Authors:** Klaus Wolkenstein, Burkhard C. Schmidt, Mathias Harzhauser

## Abstract

Polyene pigments represent a major class of pigments in present-day organisms. Their occurrence in fossils has been frequently discussed, but to date no spectroscopic evidence was found. Here, we use *in situ* Raman microspectroscopy to examine the chemistry of exceptionally well-preserved gastropod shells with colour preservation from the Middle Miocene of the Vienna Basin (Austria, Hungary). Raman signals indicative for the presence of intact, i.e. non-hydrogenated polyene pigments were obtained from the fossil gastropods, thus revealing the first record of intact polyenes in fossils. The observed Raman values are in good agreement with those of unmethylated (non-carotenoid) polyenes. Fossil polyene pigments were detected in representatives of the superfamily Cerithioidea, but not in representatives of other gastropod families with colour preservation found at the same localities. Our results show that Raman spectroscopy represents a valuable tool for the non-destructive screening of rare fossils with colour preservation for the occurrence of polyene pigments.

## Introduction

Polyene pigments are widely distributed in the three domains of life and are responsible for most yellow, orange and red colours observed in nature (Yabuzaki, 2017; Maia et al., 2021). The dominant group among the polyene pigments are the carotenoids, but non-carotenoid polyenes such as the psittacofulvins have increasingly become the subject of recent research (Maia et al., 2021). The occurrence of polyene pigments in fossils has been discussed especially in the context of molluscan shell colour (Williams, 2017; Wolkenstein, 2022), plumage pigments (Thomas et al., 2014a; Thomas et al., 2014b) and recent paleocolour reconstructions (McNamara et al., 2016; Roy et al., 2020; Vinther, 2020). However, because of the known susceptibility of polyenes to oxidation (Britton et al., 2008), the preservation potential of intact polyenes in sediments and fossils is generally considered to be low (Sinninghe Damsté and Koopmans, 1997; Williams, 2017), and to date no spectroscopic evidence for the preservation of intact polyene pigments in fossils has been reported.

Colour pattern preservation can be commonly found in Cenozoic molluscan shells (Hollingworth and Barker, 1991). Many shell colour patterns that have faded to some extent have also been revealed using UV light (Caze et al., 2010; Caze et al., 2011; Schneider et al., 2013; Hendricks, 2015 and references therein). By contrast, obvious colour preservation, i.e. preservation of distinct colours in fossils is a rare exception. An outstanding example is the small gastropod *Pithocerithium rubiginosum* (Cerithiidae), representing one of the most abundant gastropod species of the Sarmatian (Middle Miocene) of the Central Paratethys (Harzhauser and Piller, 2010). Many specimens of *P. rubiginosum* show a distinct red colouration of the beads of the spiral cords (Fig. 1A). This red colouration is well known since the 19th century (Hörnes, 1848) and it is so conspicuous that the species even was named for it (from the Latin for ‘rusty red ‘) by Eichwald (1830; 1853). In addition to *P. rubiginosum*, colour preservation can be found in a number of further gastropods from the Vienna Basin. However, despite the often distinct colouration, until now no attempts were undertaken to chemically analyse the pigments of the fossil gastropods.

**Figure 1.**
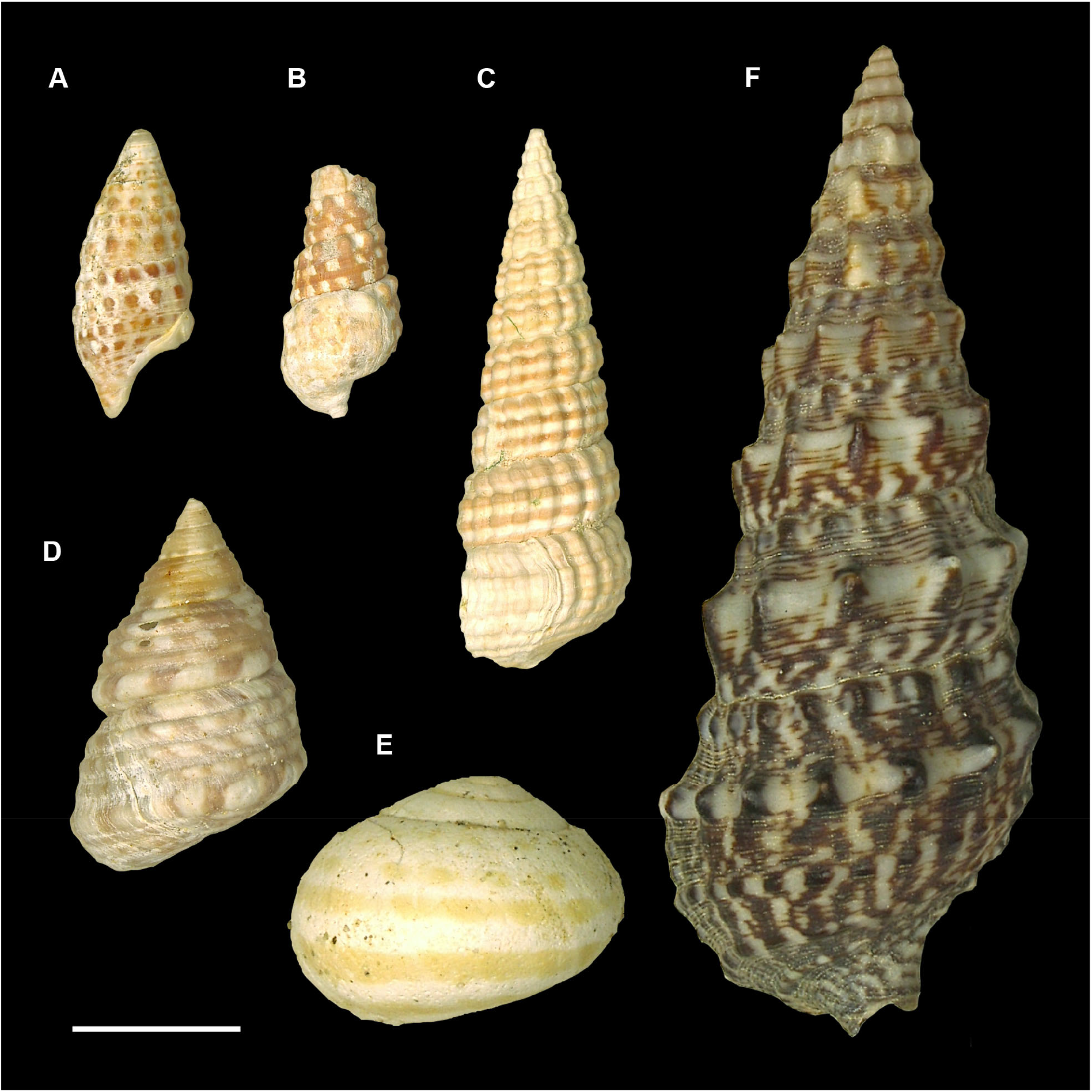
Examples of coloured Miocene and modern gastropods used for Raman measurements. A) *Pithocerithium rubiginosum*, Miocene, Nexing, Austria, NHMW 2023/0101/0001, B) *Tiaracerithium pictum*, Miocene, St. Margarethen, Austria, NHMW 2023/0103/0001, C) *Potamides disjunctus*, Miocene, Fertörákos, Hungary, NHMW 2023/0104/0002, D) *Sarmatigibbula podolica*, Miocene, Fertörákos, Hungary, NHMW 2023/0104/0003, E) *Megalotachea sylvestrina*, Miocene, Nexing, Austria, NHMW 2023/0101/0003, F) *Thericium vulgatum*, modern, Mediterranean Sea, Malacological Collection of NHMW, NHMW 2023/0105/0001. Scale bar represents 1 cm.

A survey of the literature shows that even the pigments of present-day molluscs are merely known (Williams, 2017). Some results concerning the pigments of present-day molluscs were obtained by Raman analysis. Although Raman spectroscopy cannot be considered as a method for structure elucidation in the narrower sense, it provides valuable information on the molecular framework and the nature of bonding of unknown compounds. *In situ* Raman spectroscopy suggested that polyene pigments are distributed in many present-day molluscs (Barnard and de Waal, 2006; Hedegaard et al., 2006; Ishikawa et al., 2019; Wade et al., 2019). Polyene pigments are polyunsaturated organic compounds that contain at least two conjugated carbon double bonds and include natural products such as isoprenoid carotenoids and unmethylated polyenes such as the psittacofulvins (Maia et al., 2021). Based on Raman data, those polyenes found in modern molluscs generally have been assigned as unmethylated polyenes (Barnard and de Waal, 2006; Hedegaard et al., 2006; Ishikawa et al., 2019).

In recent years, Raman spectroscopy has gained increasing popularity in the analysis of fossil samples, because it is a non-destructive method and measurements can be easily performed. On the other hand, some limitations and pitfalls have to be considered with the Raman analysis of fossil samples. In particular, the high autofluorescence of many fossils represents a serious problem, often making analysis impossible (Olcott Marshall and Marshall, 2015; Geisler and Menneken, 2021). In addition to inorganic compounds, Raman has been successfully applied to fossil organic materials such as amber (Winkler et al., 2001) and fossilized latex (Lönartz et al., 2023), although it should be considered that not all organic molecules are equally suitable for the analysis with Raman. A limitation for organic compounds is that especially in thermally more mature samples commonly only D and G bands of carbon are obtained (Peteya et al., 2017; Wolkenstein, 2022). Moreover, in some recent Raman studies on fossil samples (e.g., Wiemann et al., 2018; McCoy et al., 2020) numerous quasi-periodic signals have been obtained over the full spectral range, which, however, are supposed to represent instrumental artefacts generated by interferences from intense background fluorescence at the edge filters (Alleon et al., 2021).

In order to explore the chemical nature of colourful fossil gastropod shells from the Vienna Basin as non-destructive as possible, we decided to investigate a set of representative specimens with Raman spectroscopy. Raman appears particularly suitable for the analysis of potential fossil polyene pigments, since polyenes are known to be strong Raman scatterers (Maia et al., 2021), thus allowing for the detection of even small amounts of pigments.

## Material and Methods

A number of common gastropods showing colour preservation has been selected from the Middle Miocene (Sarmatian) of the Vienna Basin comprising specimens from four different localities: Nexing, Austria (N 48.493324°, E 16.658877°), Hautzendorf, Austria (N 48.453997°, E 16.503818°), St. Margarethen, Austria (N 47.762537°, E 16.629692°) and Fertőrákos Piusz-Puszta section, Hungary (N 47.720170°, E 16.650400°) (Fig. S1). The species represent five different genera and families (Harzhauser et al., 2023). Four of the species dwelled in coastal marine environments and one species was terrestrial. The shells were found in mixed carbonatic siliclastic sediments, ranging from sand and oolithic sand to shell coquinas.

Cerithiidae: *Pithocerithium rubiginosum* from Nexing (NHMW 2023/0101/0001) and Hautzendorf (NHMW 2023/0102/0001); paleoenvironment: coastal marine.

Batillariidae: *Tiaracerithium pictum* from Nexing (NHMW 2023/0101/0002), Hautzendorf (NHMW 2023/0102/0002), St. Margarethen (NHMW 2023/0103/0001) and Fertörákos (Hungary) (NHMW 2023/0104/0001); paleoenvironment: coastal marine.

Potamididae: *Potamides disjunctus* from St. Margarethen (NHMW 2023/0103/0002) and Fertörákos (NHMW 2023/0104/0002) ; paleoenvironment: coastal marine.

Trochidae: *Sarmatigibbula podolica* from Fertörákos (NHMW 2023/0104/0003); paleoenvironment: coastal marine.

Helicidae: *Megalotachea sylvestrina* from Nexing (NHMW 2023/0101/0003); paleoenvironment: terrestrial.

For comparison we used a specimen of the modern Cerithiidae *Thericium vulgatum* without ostracum from the Mediterranean Sea (Malacological Collection of NHMW, NHMW 2023/0105/0001). All specimens used in this study are stored in the collection of the Natural History Museum Vienna.

Samples were cleaned with acetone and coloured as well as non-coloured areas of the shell were subjected to *in situ* Raman spectroscopy. Raman spectra were acquired using a Horiba Labram HR800 UV instrument equipped with an Olympus BX41 microscope. The excitation wavelength was the 488 nm line of a diode laser with a reduced laser power (10%) of about 0.45 mW at the sample surface. The use of a 100× objective for focusing the laser on the sample surface and a confocal hole of 100 µm yielded a spatial resolution of about 1 µm lateral and 5 µm in depth. The scattered light was dispersed by a grating with 600 lines mm^−1^ and detected with a charge-coupled device (CCD) detector with 1024 × 256 pixels. Raman spectra were acquired in the range of 100–2100 cm^−1^ and 100–4200 cm^−1^ with a spectral dispersion of about 2 cm^−1^ and final spectra were averaged from 16 spectra with 15 s exposure time each (with exception of the spectrum of *Tiaracerithium* from the Miocene of St.

Margarethen that was averaged from 24 spectra with 10 s exposure time, because of the high fluorescence of the sample). The spectrometer was calibrated against the Si band at 520.4 cm^− 1^. During the measurement, the Horiba intensity correction system (ICS) was applied in order to minimize instrument specific interference effects of the edge filters that potentially may lead to quasi-periodic instrumental artefacts (Alleon et al., 2021). After measurement, adaptive background correction was applied using SpectraGryph 1.2 (coarseness: 10%, offset: 0). Uncorrected Raman spectra that show the elevated fluorescence background especially of the fossil samples are shown in the Supporting Information (Fig. S2, Fig. S3).

UV light-induced autofluorescence of samples was documented using a Canon PowerShot A700 digital camera and a Benda NU-8 KL 8W UV lamp (366 nm), with a distance of the light source to the fossils of about 10 cm. The exposure time for all UV photographs was two seconds.

A further specimen of *Pithocerithium rubiginosum* from Nexing with preservation of red dots (see Fig. 2E) was ground to a fine powder and stirred overnight in EDTA solution (pH 8.0) in order to dissolve the shell without destroying potential polyene pigments (which might be degraded by acids). Following centrifugation, the supernatant was removed and the orange-brown residue was extracted with distilled water. The yellowish solution was subjected to solid-phase extraction. The sorbent (Bondesil C18, 40 µm) was conditioned with acetonitrile, followed by water. The sample solution then was loaded onto the column, and the sorbent was washed with water. The yellow pigments were eluted with acetonitrile and the solvent was removed by evaporation under a stream of nitrogen at 40°C. Pigments were transferred to a CaF_2_ disk. Raman spectra were acquired as described above, however, with a strongly reduced laser power (1%) of about 0.045 mW at the sample surface to avoid any deterioration of the organic sample.

**Figure 2.**
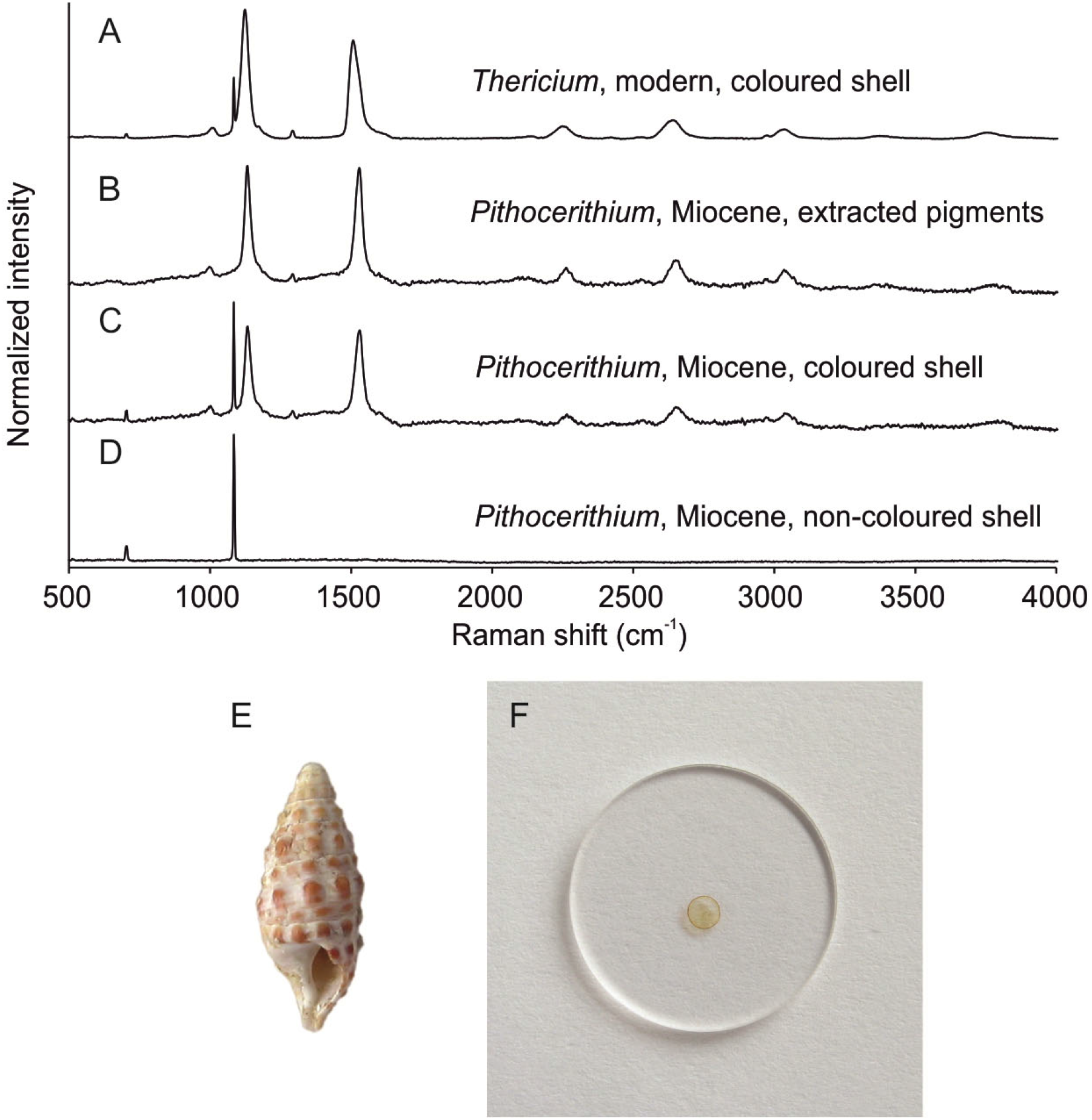
Raman analysis of shell pigments of the Miocene gastropod *Pithocerithium* and the modern gastropod *Thericium*. A) *In situ* Raman spectrum of the brown coloured shell of the modern *Thericium vulgatum* (see Fig. 1F), B) Raman spectrum of the isolated orange polyene pigment of *Pithocerithium rubiginosum* from the Miocene of Nexing (see F), C) *In situ* Raman spectrum of the red coloured shell of *Pithocerithium rubiginosum* from the Miocene of Nexing (see Fig. 1A), D) *In situ* Raman spectrum of the non-coloured shell of *Pithocerithium rubiginosum* from the Miocene of Nexing (see Fig. 1A). E) Specimen of *Pithocerithium rubiginosum* from the Miocene of Nexing used for extraction of pigments (height is 1.5 cm). F) Isolated orange polyene pigment of *Pithocerithium rubiginosum* (see E) on CaF_2_ disk (diameter of disk is 2 cm). Raman spectra are background corrected and normalized. The signal at 1085 cm^−1^ is due to carbonate.

## Results

The majority of specimens of *Pithocerithium rubiginosum* found at the localities Nexing and Hautzendorf (Fig. S1) show a distinct orange to red colouration of the beads of the spiral cords (Fig. 1A). Orange to red colouration can also be observed in many shells of *Tiaracerithium pictum* from all four investigated localities (Fig. 1B). In the shells of *Potamides disjunctus* from St. Margarethen and Fertörákos, spiral lines with a tan to reddish colour can be found (Fig. 1C), although in general the colouration in *Potamides* is less distinct than in *Pithocerithium* and *Tiaracerithium*. In contrast to *Granulolabium* and *Potamides*, specimens of *Sarmatigibbula podolica* with colour preservation from Fertörákos show only a tan colour (Fig. 1D). Specimens of *Megalotachea sylvestrina* from Nexing show yellow spiral bands (Fig. 1E).

Raman spectra indicate preservation of the original aragonite shell (several signals between 125 und 225 cm^−1^, but no signal at about 280 cm^−1^ that would be indicative for calcite) in all investigated gastropod specimens (Fig. S2, Fig. S3). Further signals characteristic for carbonates occur at 704 and 1085 cm^−1^, the latter being the most intense carbonate signal (Fig. 2, Fig. S2, Fig. S3). From the Raman spectra, it can also be excluded that the reddish colouration of the shells is due to iron oxides such as hematite, because no corresponding signals (ca. 225, 290, 410, 610, 1320 cm^−1^) (de Faria et al., 1997; Marshall and Olcott Marshall, 2011) were found.

Whereas no signals in addition to the carbonate bands were obtained from non-coloured shell areas of the gastropods (Fig. 2D), Raman spectra of coloured areas of the fossil gastropods *Pithocerithium, Tiaracerithium* and *Potamides* from different localities show distinct signals at about 1100 and 1500 cm^−1^ (Fig. 2C, Fig. S2, Table 1), different than common signals of D and G bands (at about 1350 and 1600 cm^−1^). Both orange and red colouration show the same Raman signals, differing only in the intensity of the signals. This indicates that only the concentration of pigments is different in the individual shells. In those shells with the most intense colouration (*Pithocerithium* and *Granulolabium*) further signals at about 1000, 1300, 2200, 2600 and 3000 cm^−1^ (Fig. 2C, Table 1) can be observed. All these signals are characteristic for polyene pigments (Barnard and de Waal, 2006; Hedegaard et al., 2006) and can be interpreted corresponding to known Raman bands of polyenes. The signals of highest intensity at about 1500 cm^−1^ (ν_1_) and at about 1100 (ν_2_) can be assigned to C=C and C–C stretching bands respectively and the weaker signals at about 1300 cm^−1^ (ν_3_) and at about 1000 (ν_4_) to CH=CH and CH bending bands (Barnard and de Waal, 2006). The signals above 2000 cm^−1^ represent overtone and combination bands (mainly of the intense ν_1_ and ν_2_ bands). The specific polyene signals are also observed in the Raman spectrum of a dark brown coloured present-day *Thericium vulgatum* (Fig. 2A). The ν_1_ signals of the red coloured *Pithocerithium* are shifted by about 20 cm^−1^ to higher wavenumbers compared to the brown coloured *Thericium* (Table 1). This shift can be explained by the conjugation length of the polyene backbones, which is higher for brown polyenes than for red polyenes (Ishikawa et al., 2019).

**Table 1.** Raman signals (cm^−1^) of shell pigments of Miocene gastropods compared to those of the modern gastropod *Thericium* n.d., not determined. Because of the compared to other samples lower signal intensity, for these samples Raman spectra were acquired only in the range of 100–2100 cm^−1^

| Gastropod | Superfamily | Locality | $\nu_4$ | $\nu_2$ | $\nu_3$ | $\nu_1$ | $2 \nu_2$ | $\nu_1 + \nu_2$ | $2 \nu_1$ |
| --- | --- | --- | --- | --- | --- | --- | --- | --- | --- |
| *Pithocerithium* | Cerithioidea | Nexing | 1002 | 1133 | 1293 | 1530 | 2263 | 2654 | 3038 |
| *Pithocerithium* (extract) | Cerithioidea | Nexing | 999 | 1132 | 1293 | 1529 | 2263 | 2652 | 3035 |
| *Pithocerithium* | Cerithioidea | Hautzendorf | 997 | 1133 | 1295 | 1530 | 2266 | 2653 | 3037 |
| *Tiaracerithium* | Cerithioidea | Nexing | 999 | 1134 | 1293 | 1531 | 2267 | 2653 | 3038 |
| *Tiaracerithium* | Cerithioidea | Hautzendorf | 1000 | 1134 | 1292 | 1530 | n.d. | n.d. | n.d. |
| *Tiaracerithium* | Cerithioidea | St. Margarethen | 1002 | 1134 | 1293 | 1531 | n.d. | n.d. | n.d. |
| *Tiaracerithium* | Cerithioidea | Fertőrákos | 1001 | 1134 | 1296 | 1531 | n.d. | n.d. | n.d. |
| *Potamides* | Cerithioidea | St. Margarethen | 1002 | 1134 | 1296 | 1531 | n.d. | n.d. | n.d. |
| *Potamides* | Cerithioidea | Fertőrákos | 1001 | 1134 | 1296 | 1531 | n.d. | n.d. | n.d. |
| *Sarmatigibbula* | Trochoidea | Fertőrákos | – | – | – | – | n.d. | n.d. | n.d. |
| *Megalotachea* | Helicoidea | Nexing | – | – | – | – | n.d. | n.d. | n.d. |
| *Theridium* | Cerithioidea | Mediterranean Sea | 1009 | 1124 | 1293 | 1507 | 2249 | 2639 | 3037 |

Remarkably, taxonomic differences in the composition of pigments are still recorded in the gastropod shells. Although evidence for polyene pigments was found in gastropods from all investigated localities in the Vienna Basin, not all species with preservation of colour patterns from Nexing (e.g. *Megalotachea sylvestrina*) show the presence of polyene pigments (Fig. S3, Table 1) (intense signals at 1100 und 1500 cm^−1^ are missing). No polyene pigments were also found in *Sarmatigibbula* with colour pattern preservation from Fertörákos (Fig. S3, Table 1). The presence or lack of polyenes in the gastropod shells (fossil and modern) is also in agreement with the fluorescence properties of the shells as observed under UV light. Whereas gastropods that yielded Raman signals characteristic for polyenes show no fluorescence of the pigments, the colour patterns of *Megalotachea* and *Sarmatigibbula* show distinct yellow or red fluorescence, respectively (Fig. S4), suggesting the presence of other pigments than polyenes.

Following dissolution of the shell, the pigments of a specimen of *Pithocerithium rubiginosum* with preservation of red dots were extracted and isolated (Fig. 2E). Raman analysis of the isolated orange pigments (note that the carbonate signal at 1085 cm^−1^ has disappeared) yielded all characteristic polyene signals (Fig. 2B, Table 1) that were previously observed in the *in situ* Raman spectrum of the pigmented gastropod shell (Fig. 2C, Table 1) with even better signal-to-noise ratio, indicating that the Raman signals are due to the orange pigments.

## Discussion

The specific Raman signals of the fossil gastropod pigments show the presence of C-C and C=C bonds, which have to be arranged at least to some extent in the form of conjugated double bonds, as obvious from the orange to red colour of the compounds, thus indicating the presence of original, still intact (non-hydrogenated) polyene pigments. The Raman signals of the pigments of the fossil Cerithioidea are very similar to those obtained from the pigments of the modern representative *Thericium* and to those of many other modern molluscan pigments that have been attributed to polyenes. However, to the best of our knowledge such Raman signals have not yet been reported from the fossil record. Raman spectroscopy has been used to search for carotenoid pigments in amber-preserved feathers, but distinctive carotenoid-informative bands (at 1530 and 1153 cm^−1^) have been only obtained for present-day, but not for fossil feathers (Thomas et al., 2014b). Putative carotenoproteins were reported for the brachiopod *Calloria* from the Late Pleistocene of New Zealand (Curry, 1999) and the gastropod *Ecphora* from the Middle Miocene of North America (Nance et al., 2015), but both studies mainly focused on the characterization of the proteinaceous material or the content of amino acids with very little information on the associated pigments. Otherwise, only hydrogenated carotenoids (non-coloured carotanes) have been reported from fossil organisms, such as *β*-carotane that was detected in *Pseudofagus* leaves from the Miocene of Clarkia, Idaho (Giannasi and Niklas, 1981).

The discovery of still intact polyene pigments in gastropod shells of Middle Miocene (Sarmatian) age is highly surprising, since the preservation potential of unsaturated polyene pigments such as carotenes in contrast to hydrogenated carotenoids is very low (Sinninghe Damsté and Koopmans, 1997). It should be also noted that no indications for the presence of hydrogenated polyenes (no signal at 1455 cm^−1^) can be found in the Raman spectra of the fossil gastropods (Marshall and Olcott Marshall, 2010). The previously oldest intact polyenes, all of them found in sediments, not in macrofossils, were the carotenoid isorenieratene from a Late Miocene (Messinian) marl from Italy (Keely et al., 1995) and an unspecified diaromatic carotenoid from a Lower Miocene clay from the Blake Bahama basin in the Western North Atlantic Ocean (Cardoso et al., 1978) (Sinninghe Damsté and Koopmans, 1997; Hopmans et al., 2005).

The exact wavenumber positions of the polyene Raman bands (in particular of the ν_1_ and ν_2_ bands) depend on the length of the polyene chain, substituents attached to the chain, its molecular configuration and potential binding to other molecules (e.g. proteins) (Schaffer et al., 1991; Hedegaard et al., 2006; Maia et al., 2021). It has been shown that the ν_2_ band can be used as a reliable indicator for the presence or absence of methyl groups in polyenes (Barnard and de Waal, 2006; Maia et al., 2021). Whereas methylated polyenes such as carotenoids have ν_2_ values of about 1150 to 1158 cm^−1^ (Barnard and de Waal, 2006), those of unmethylated polyenes such as polyanes (psittacofulvins) have ν_2_ values that are about 20 to 30 cm^−1^ lower than those of the methylated polyenes (Maia et al., 2021). Therefore, the ν_2_ values observed in the present work (1132 to 1134 cm^−1^) (Table 1) rather points to the occurrence of unmethylated polyenes in fossil and modern representatives of the Cerithioidea. This is also in agreement with a generally observed wide occurrence of unmethylated polyenes in modern molluscs (Barnard and de Waal, 2006; Hedegaard et al., 2006; Ishikawa et al., 2019).

Furthermore, it has been demonstrated that with increasing length of the conjugated chain in unmethylated polyenes the wavenumber position of the ν_1_ band decreases (Schaffer et al., 1991). These data obtained on synthetic polyenes have been successfully applied to the characterization of unmethylated polyenes from differently coloured modern molluscan shells (Hedegaard et al., 2006; Ishikawa et al., 2019). By comparison of the Raman peak maxima measured in the present work with those of the synthetic polyenes (Schaffer et al., 1991), the number of conjugated double bonds of the polyenes in the fossil cerithioidean gastropods and the modern *Thericium* can be estimated to be n = 9 and n = 12, respectively, corresponding very well to the observed orange-red and brown colour of the gastropod shells. It is also remarkable that the peaks in *Thericium* are broader than in the fossil representatives, suggesting a broader range of conjugated double bonds in the pigments of the modern gastropod.

It is well known that polyene pigments such as carotenoids in nature often occur as carotenoproteins, which have different colours than the free carotenoids (Britton et al., 2008). Interaction of unmethylated polyenes (psittacofulvins) with the protein keratin have been reported too (Stradi et al., 2001). However, although proteinaceous material has been reported from shells of the Miocene gastropod *Ecphora* (Nance et al., 2015), in the present work, no indications were found that the polyenes of the cerithioidean gastropods could be bound to proteins. Nevertheless, it has to be considered that the actual colour of the polyene-containing Miocene gastropods during lifetime may have been different than orange-red, if they were bound to proteins.

A prerequisite for the exceptional colour preservation in the investigated fossil gastropods is certainly the general low thermal maturity of organic material from the Neogene sediments in the Vienna Basin (Sachsenhofer, 1992). A low degree of diagenetic changes is also indicated by the preservation of the original aragonite shell. Furthermore, the occurrence of the pigments within the carbonate matrix of the gastropod shells has obviously contributed to the exceptional preservation.

It is very interesting that reddish colouration of shells and corresponding Raman signatures for intact polyenes were only found in fossil members of the superfamily Cerithioidea (*Pithocerithium, Potamides, Tiaracerithium*), but not in members of other gastropod families with colour preservation from the same localities such as *Sarmatigibbula* (superfamily Trochoidea) and *Megalotachea* (superfamily Helicoidea) (Fig. 1, Table 1). This indicates that the occurrence of polyene pigments in ancient gastropods is related to certain taxonomic groups.

## Conclusions

Important information on the chemistry of organic compounds in fossils can be obtained from Raman bands. Whereas typically D and G bands are obtained from thermally mature material, more characteristic bands can be obtained from thermally immature samples. *In situ* Raman spectroscopy indicates the preservation of intact polyene pigments in about 12 million-year-old orange to red coloured gastropod shells from the Middle Miocene of the Vienna Basin. As far as known, these pigments represent the oldest record of intact polyene pigments in fossils. Moreover, the preservation of these pigments is observed especially within Cerithioidea, but not in two other tested gastropod superfamilies. The results show that *in situ* Raman analysis enables the non-destructive analysis of rare fossils with colour preservation and thus provides a valuable tool for the screening of further fossils on the occurrence of polyene pigments.

## Acknowledgements

We thank Susanne Affenzeller for bringing coloured specimens of *M. sylvestrina* and *P. rubiginosum* from the Vienna Basin to the attention of K.W. This work was supported by the Deutsche Forschungsgemeinschaft (WO 1491/5-1).

