## Supporting Information for "Polyene-based colouration preserved in 12 million-year-old gastropod shells"

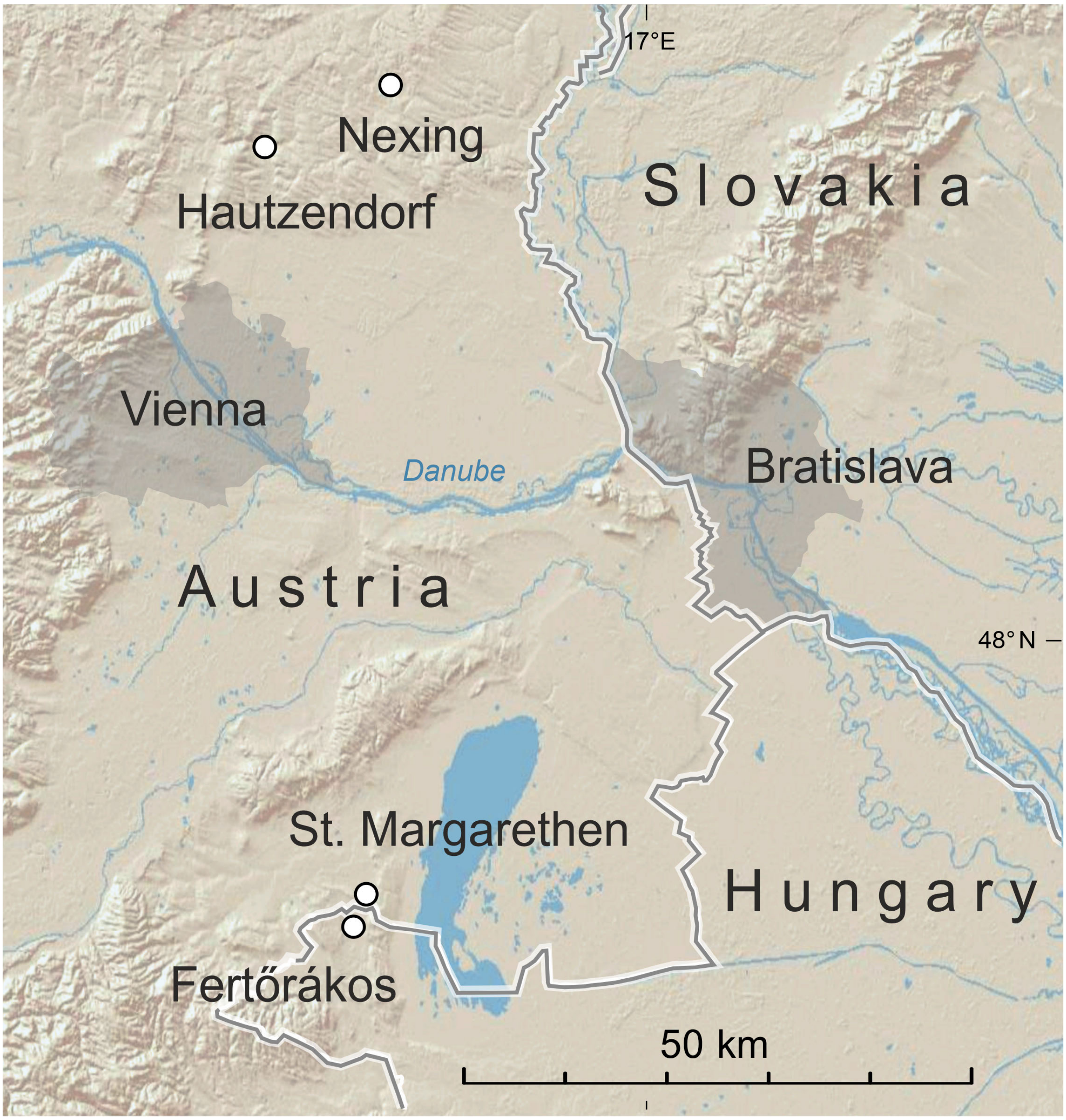


**Figure S1.** Overview of the Vienna Basin with localities of studied fossil gastropods with colour preservation. Basemap created with © 2009 Esri (World Shaded Relief).

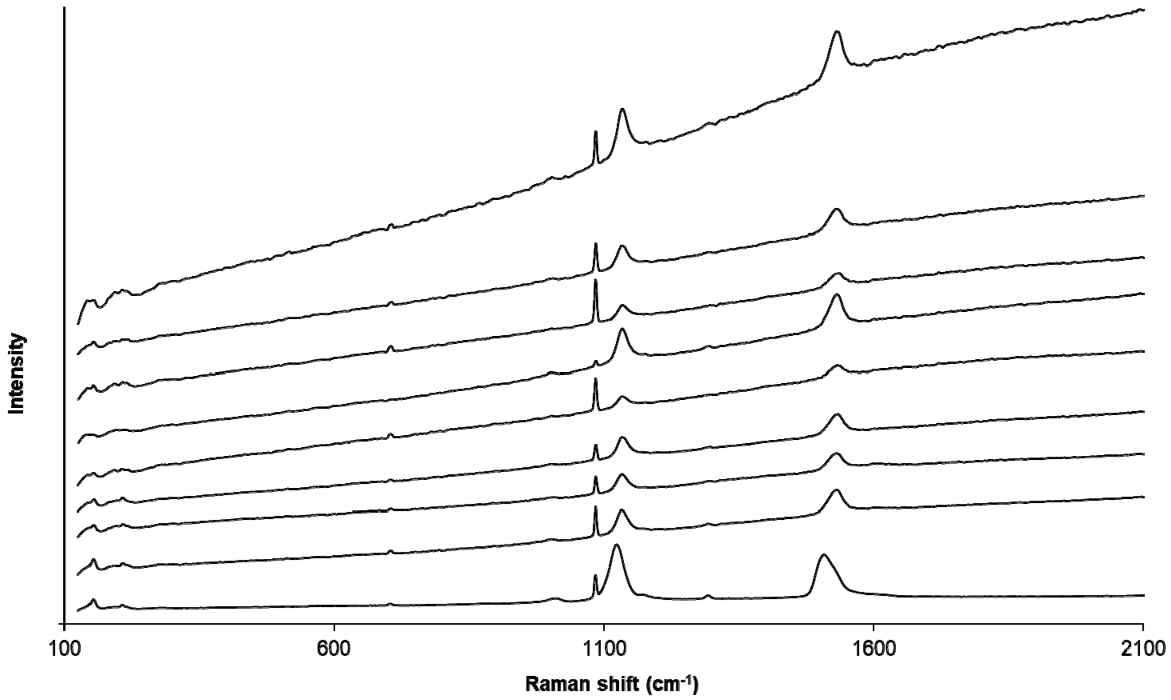


**Figure S2.** *In situ* Raman spectra (without background correction) of polyene-containing coloured shell areas. From top to bottom the spectra correspond to: *Tiaracerithium pictum*, Miocene, St. Margarethen; *Tiaracerithium pictum*, Miocene, Hautzendorf; *Tiaracerithium pictum*, Miocene, Fertörákos; *Tiaracerithium pictum*, Miocence, Nexing; *Potamides disjunctus*, Miocene, St. Margarethen; *Potamides disjunctus*, Miocene, Fertörákos; *Pithocerithium rubiginosum*, Miocene, Hautzendorf; *Pithocerithium rubiginosum*, Miocene, Nexing; *Thericium vulgatum*, modern, Mediterranean Sea. The signal at 1085 $cm^{-1}$ is due to carbonate. Raman spectra are vertically shifted for better visibility.

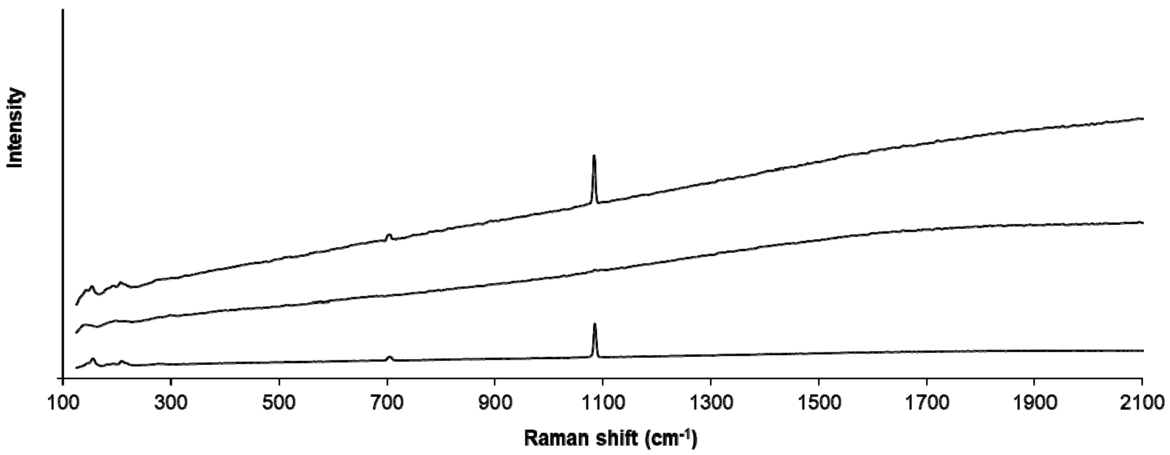


**Figure S3.** *In situ* Raman spectra (without background correction) of non-coloured shell areas and coloured shell areas that yielded no polyene signals. From top to bottom the spectra correspond to: *Sarmatigibbula podolica*, coloured shell, Miocene, Fertörákos; *Megalotachea sylvestrina*, coloured shell, Miocene, Nexing; *Pithocerithium rubiginosum*, non-coloured shell, Miocene, Nexing. Raman spectra are vertically shifted for better visibility.

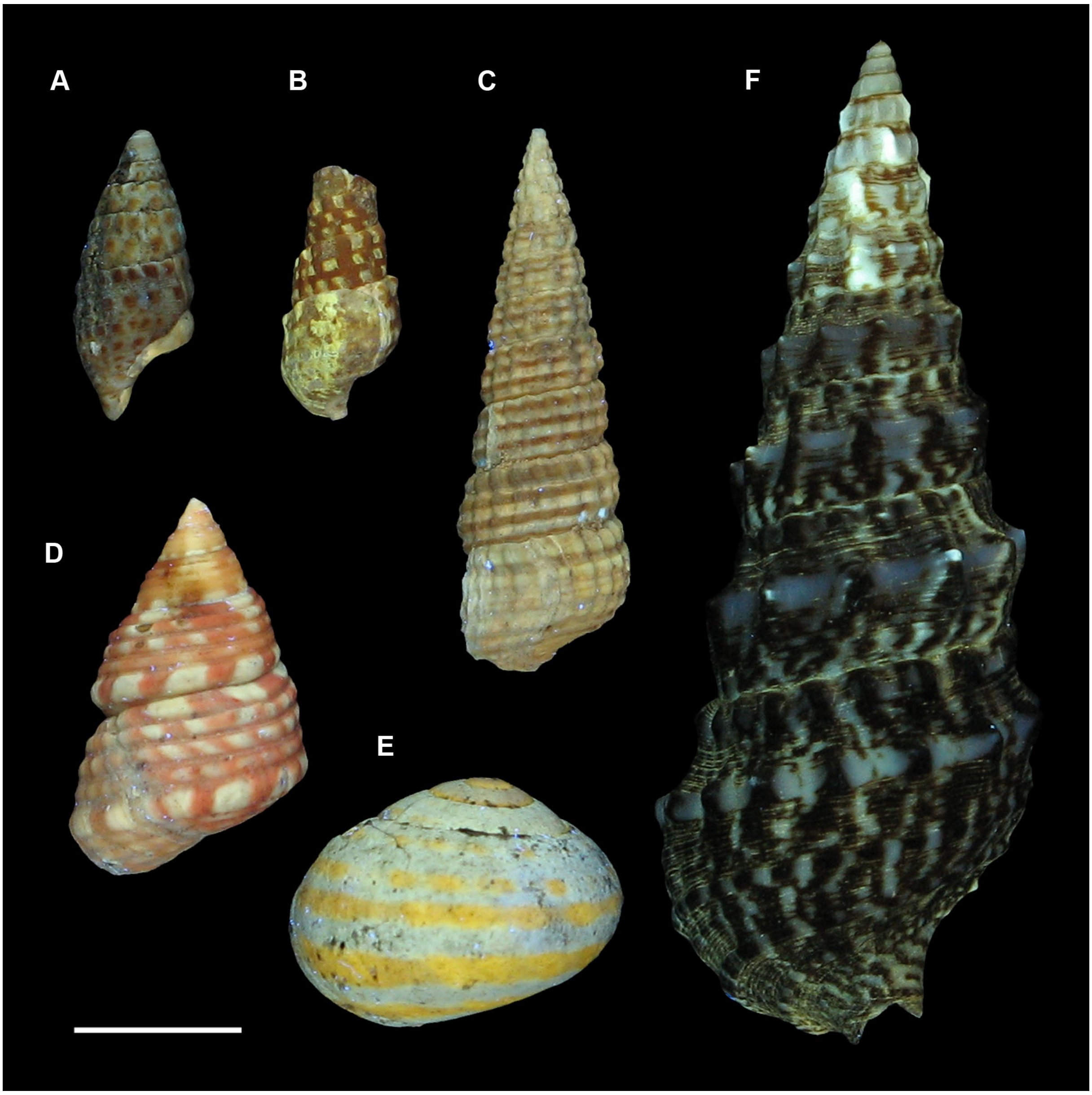


**Figure S4.** Coloured Miocene and modern gastropods under UV light. A) *Pithocerithium rubiginosum*, Miocene, Nexing, Austria, NHMW 2023/0101/0001, B) *Tiaracerithium pictum*, Miocene, St. Margarethen, Austria, NHMW 2023/0103/0001, C) *Potamides disjunctus*, Miocene, Fertörákos, Hungary, NHMW 2023/0104/0002, D) *Sarmatigibbula podolica*, Miocene, Fertörákos, Hungary, NHMW 2023/0104/0003, E) *Megalotachea sylvestrina*, Miocene, Nexing, Austria, NHMW 2023/0101/0003, F) *Thericium vulgatum*, modern, Mediterranean Sea, Malacological Collection of NHMW, NHMW 2023/0105/0001. Scale bar represents 1 cm.
